# SICD6mA: Identifying 6mA Sites using Deep Memory Network

**DOI:** 10.1101/2020.02.02.930776

**Authors:** Wenzhong Liu, Hualan Li

## Abstract

**Background:** DNA N6-methyladenine (6mA) is a kind of epigenetic modification in prokaryotes and eukaryotes, which involves multiple biological processes, such as gene regulation and tumorigenesis. Identifying 6mA contributes to understand its regulatory role. Therefore, to satisfy the needs of large-scale preliminary screening, it is necessary to develop the high-quality computational models for the rapid identification of 6mA sites. However, the existing calculation approaches are mostly specific to rice, and they have not been extensively applied to human genome.

**Results:** This study proposed a classification method of deep learning based on the memory mechanism named SICD6mA. In addition, the large benchmark datasets were constructed for human and rice, respectively, which integrated the recently reported 6mA sites. According to the evaluation results, SICD6mA displayed favorable robustness during cross-validations, which achieved the area under the curve (AUC) values of 0.9824 and 0.9903 for Human and Rice’s genomes in independent test evaluations, separately.

**Conclusions:** The successful prediction rate of 6mA sites on cross-species genomes exhibited higher accuracy than that of the state-of-the-art methods. For the convenience of experimental scientists, the user-friendly tool SICD6mA was developed to predict the cross-species 6mA sites, thereby accelerating and facilitating future cross-species genome research.

## Background

DNA N6-methyladenine(6mA) refers to methylation at the N6site of adenine[1]. Typically, DNA 6mA modification is the most common DNA modification in prokaryotic genomes[2], which is once thought not to exist in eukaryotes[3] (including humans), because it is not detected in earlier studies. With the development of high-throughput sequencing technology, numerous 6mA modifications have been detected in different multicellular eukaryotes. In recent years, 6mA modification has been found to play an important role in DNA epigenetic modification of eukaryotes. DNA methylation represents a key element in epigenetic regulation, and DNA 6mA modification exerts a vital part in regulating gene expression[4], germ cell differentiation[5] etc. It has been reported in the genomes of ***E. coli[6], Drosophila melanogaster***[7], ***rice***[8], and ***Arabidopsis***[9] etc. Currently, the 6mA sites can be detected in mouse and human cells using the sensitive detection technology. 6mA is extensively distributed throughout the human genome. Of note, [G/C]AGG[C/T] is the most frequent motif at the 6mA modification site[10]. Different types of cancer cells can be distinguished from normal cells based on the heterogeneities in methylation of the CpG islands. It is of great significance to identify the potential function and clinical value of 6mA sites in human tumorigenesis[11]. However, DNA 6mA is lowly expressed, and 6mA sites are unevenly distributed throughout the genome; particularly, the biological function of 6mA in advanced eukaryotes has not been comprehensively understood yet. To shed more light on its biological function, it is essential to characterize its position on the genome, which is related to understanding the epigenetic modification process through investigating the distribution of DNA 6mA[12].

The recent development of experimental technology is in favor of finding DNA 6mA modification sites. Specifically, the experimental techniques for identifying 6mA sites include mass spectrometry[13], immunoprecipitation[14], and next-generation sequencing[15], among which, single-molecule real-time sequencing (SMRT)[16] is the mainstream experimental technique[17]. Nonetheless, the analytical techniques are costly, complicated, and usually labor-intensive, making it difficult to identify 6mA across the genome. Additionally, the method of antibody detection is susceptible to bacterial contamination, while the SMRT sequencing technique can sometimes hardly distinguish 6mA from N1-adenine (1mA) site[18]. To this end, it is of essential necessity to develop a calculation method for the rapid identification of 6mA site.

Computational methods for predicting 6mA sites are still in their infancy[19], and a number of prediction tools have been reported so far. Notably, iDNA6mA-PseKNC[20] is the first prediction approach to predict 6mA sites in the ***Mus.musculus*** genome, and the prediction accuracy for ***Mus.musculus*** and microbe on the dataset has been verified to be quite high. i6mA-Pred[21] is the first method utilized to identify rice genome through support vector machines (SVM), which employs the chemical characteristics and frequency of nucleotides to encode DNA sequences. In addition, i6mA-Pre provides a benchmark dataset of 6mA-rice-Chen. Various prediction methods are dependent on the dataset. As for MM-6mAPred[22], the Markov model based on the 6mA-rice-Chen dataset, achieves superior prediction capability of 6mA sites to i6mA-Pred. On the other hand, SDM6A[23] is also an approach that integrates coding methods with machine learning on the basis of the 6mA-rice-Chen dataset. After training and evaluation, its performance is likewise better than those of i6mA-Pred and iDNA6mA-PseKNC. csDMA[19] implements distinct algorithms for model optimization using the datasets obtained from iDNA6mA-PseKNC and i6mA-Pred. Meanwhile, i6mA-DNCP[18]employs the dinucleotide composition and dinucleotide-based DNA characteristics to identify 6mA sites in rice genome with high sensitivity, and it represents a novel idea for extracting the sequence features of 6mA sites by *k-mer* encoding. Deep learning is a powerful approach used to predict the 6mA sites. Moreover, Lu et al. proposed another 6mA benchmark dataset for rice genome, namely, 6mA-rice-Lv[24]. iDNA6mA-rice is another calculation tool to predict 6mA sites in rice genomes, which is associated with superior prediction performance on both 6mA-rice-Lv and 6mA-riceChen datasets. At present, SNNRice6mA[25], a convolutional neural network on6mA-rice-Chen and 6mA-rice-Lv datasets, is the optimal approach.

Nonetheless, limited by the computing resources, some web servers of the above-mentioned tools have very limited capabilities to handle the large sequences of online submissions. In the meantime, it is difficult to optimize the existing models and to improve the prediction accuracy in the absence of large datasets for human and rice. Therefore, it is necessary to enhance the performance and accuracy of existing computational methods for the preliminary large-scale screening of eukaryotic genomes. To this end, this study was carried out aiming to predict the 6mA sites in human and rice genomes.

The reported experimental data allow us to build these datasets and to develop the computational models. In this paper, these large datasets of Human and Rice Nip were constructed from the recent reported 6mA sites in human and rice respectively. Moreover, a novel and effective deep learning model SICD6mA (**Figure 1**) was proposed to improve the prediction accuracy of 6mA sites, due to the great insufficiency of calculation methods for humans and plants in this field. SICD6mA directly encoded a sequence based on the serial numbers of three consecutive nucleic acids, which was deemed as the 3-*mer* ID(**Figure 1.A**). In addition, it extracted the information of interest and identified the 6mA sites by the Gated Recurrent Units (GRU) network on the basis of the memory mechanism(**Figure 1.B**). Also, the SICD6mA was validated repeatedly using datasets. According to the results, the as-proposed model SICD6mA attained very stable performance, with significantly superior prediction accuracy to the current state-of-the-art methods. Furthermore, the user-friendly tool SICD6mA was also developed based on the CPU or GPU platform for biologists, and the codes and tools were completely accessible. Prior to high-throughput screening, SICD6mA was able to quickly identify the potential 6mA sites in human and rice genomes. To sum up, SICD6mA proposed in this study is a primary tool to more accurately predict 6mA sites in human and rice genomes, which accelerates and facilitates future research on human and rice genomes.

**Figure 1.**
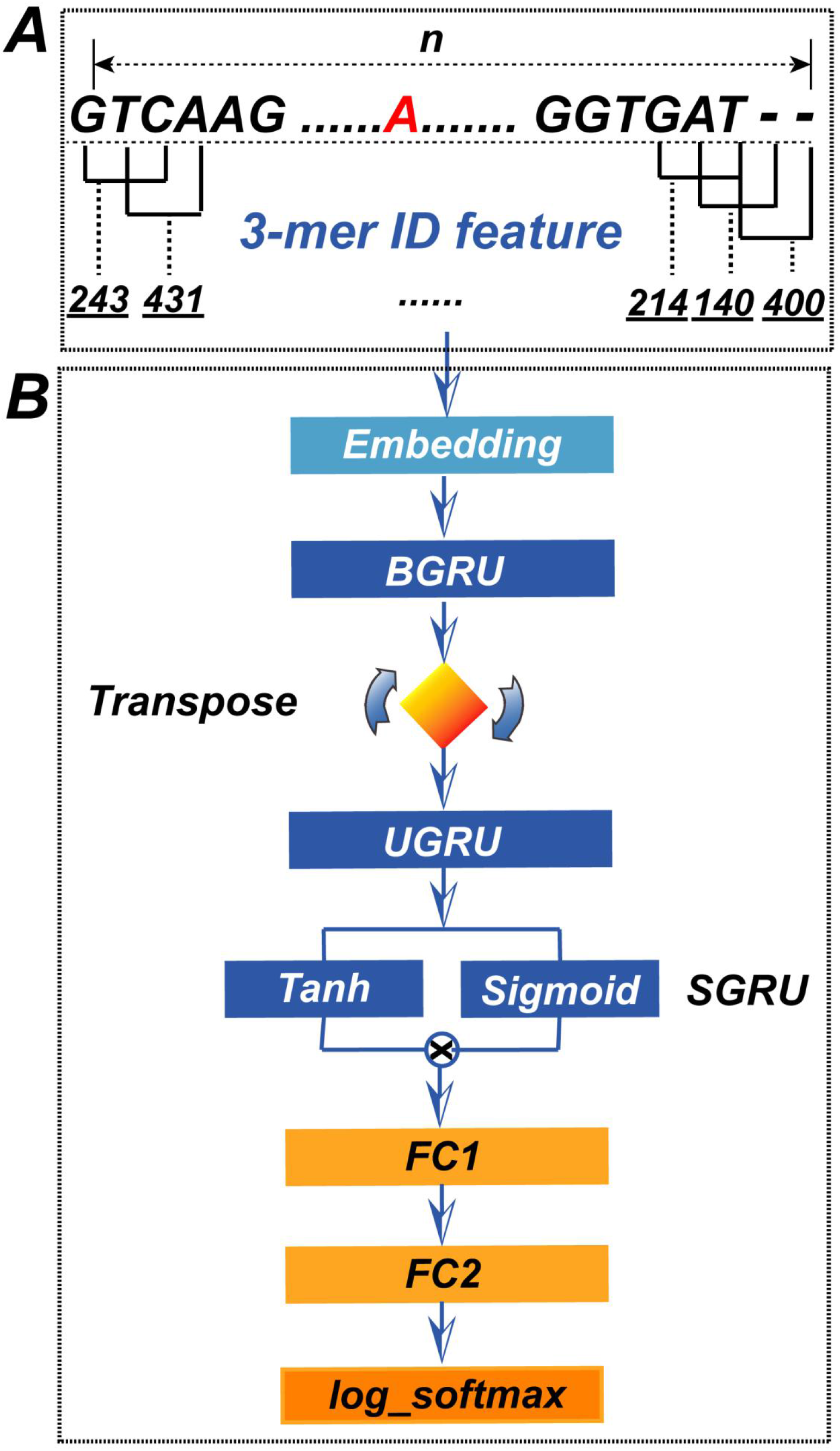
Overall Framework of SICD6mA. The two steps include: **A**. sequence encoding using *3-mer* ID feature. **B**. feature extracting and classification scoring by GRU networks based on memory mechanism.

## Results

### Overall framework of SICD6mA

In this study, the SICD6mA model based on GRU networks and memory mechanism was constructed to deal with the *3-Mer* ID feature and to identify the 6mA sites (**Figure 1**). The SICD6mA model contains eight stacked components, among which, the input vector in the first layer of SICD6mA is the *3-Mer* ID feature, an integer vector, the sort numbers of three consecutive nucleic acids on a sequence (**Figure 1. A**). A word-vector embedding layer converts the vector into a two-dimensional float matrix, namely, the embedding features (**Figure 1.B**). The Bi-directional Gated Recurrent Units (BGRU) layer learns the embedding features and extracts the first level feature map. Thereafter, SICD6mA utilizes the transpose operation to rotate the feature map particularly. Meanwhile, the non Bi-directional Gated Recurrent Units (UGRU) layer extracts the instantly second level feature map from the rotated map; subsequently, the Simple Gated Recurrent Units (SGRU) layer extracts the third level abstract features. Finally, the fully connected layers FC1 and FC2 classify the abstract features, while the log_softmax layer generates the score based on the classified features.

### Cross-validation evaluation on performance

To evaluate the robustness and sensitivity of SICMD6mA, the 5-fold cross-validation was performed based on the training sets that were comprised of both Human and Rice Nip sets. The redundant sequences of each dataset were removed, which avoided overestimation or underestimation resulted from the homologous sequences. In the 5-fold cross-validation, the training set was randomly divided into 5 groups with the same size. In each step, one group of prediction was invoked as the test set, while the remaining 4 were retained as the training set. Finally, the prediction score of 5 test sets was integrated into one file to draw the receiver operating characteristic (ROC) curve and to calculate the total area under the curve (AUC).

According to **Figure 2.A**, the AUC value of SICD6mA was as high as 0.9813 in human datasets, while that was 0.9897 in the Rice Nip dataset. Besides, the ROC curves of SICD6mA showed no great fluctuations and remained stable across different species, revealing the reliability and high prediction accuracy of the method. Interestingly, the AUC values of SICD6mA in the two datasets were quite high, which indicated that SICD6mA was adequately trained in the large datasets. **Figure 2.B** displays the precision-recall (PR) curves of the training sets. Clearly, the AP values of SICD6mA in human and Rice Nip datasets were 0.9822 and 0.9881, respectively. The two large datasets were balanced without redundancy accordingly. The two PR curves were also balanced with no sharp fluctuation, suggesting that SICD6mA attained the highest accuracy and recall, with consistent stability between the two species datasets.

**Figure 2.**
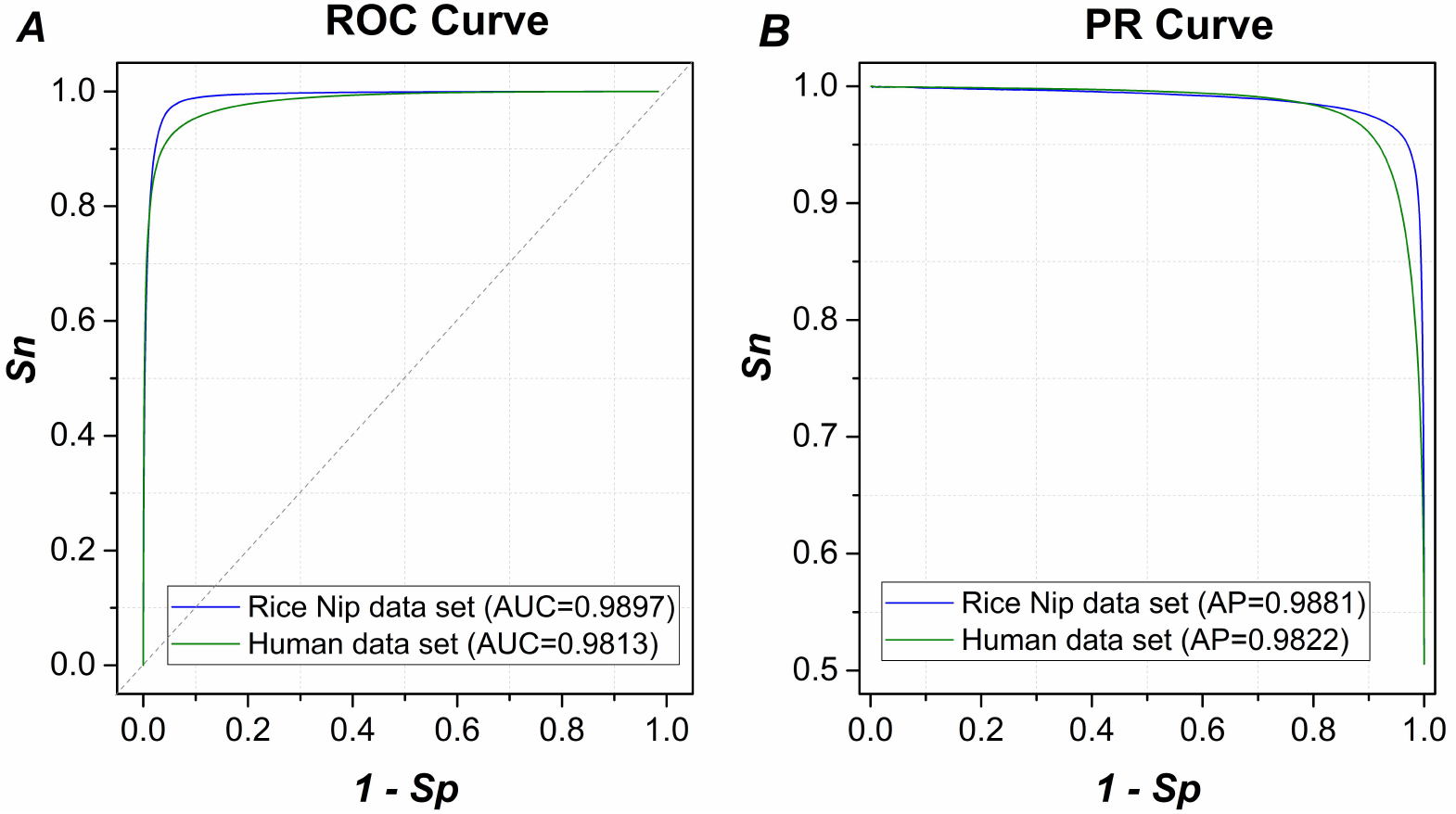
Performance evaluation based on training sets. **A.** Receiver Operating Characteristic (ROC) curves between Human and Rice Nip data sets. **B.** Precision-Recall (PR) curves between Human and Rice Nip data sets.

### Comparison of performance on independent test sets

Further, the sensitivity and accuracy of the prediction model were also evaluated based on the independent test sets. The deep learning or machine learning approaches, including i6mA-DNCP, MM-6mAPred, SNNRice6mA and csDMA have been reported recently, which achieved more accurate prediction results. In this study, the performance of SICD6mA was compared with the aforementioned methods thanks to the open source codes. To be specific, each method was trained on the training datasets and evaluated on the independent test sets. Finally, the ROC curves and PR curves of these methods were plotted according to the prediction scores of independent test sets, and then the AUC values, AP values and evaluation metrics were calculated accordingly.

The model prediction accuracy might be reliably improved due to the use of large datasets, and the prediction performance of SICD6mA was also more stable. To prove this, ROC curves were plotted and the AUC values were calculated based on the prediction results of independent test evaluations. **Figure 3.A** and **Figure 3.B** shows the ROC curves of several methods obtained by independent test evaluations. Obviously, the ROC curves of SICD6mA were on the top of other methods in the two figures. As for the Human dataset, the AUC values obtained by SICD6MA, SNNRice6mA, i6mADNCP, csDMA, and MM-6mAPred were 0.9824, 0.9748, 0.9730, 0.9719 and 0.8726, respectively; apparently, SICD6mA attained the highest AUC (**Figure 3.A**). With regard to the Rice Nip dataset, the AUC values acquired by SICD6MA, SNNRice6mA, i6mADNCP, csDMA, and MM-6mAPred were 0.9903, 0.9856, 0.9851, 0.9843, and 0.9456, respectively; clearly, the AUC of SICD6mA was higher than those of other methods (**Figure 3.B**). Overall, SICD6mA achieved quite high prediction performances in two genomes.

**Figure 3.**
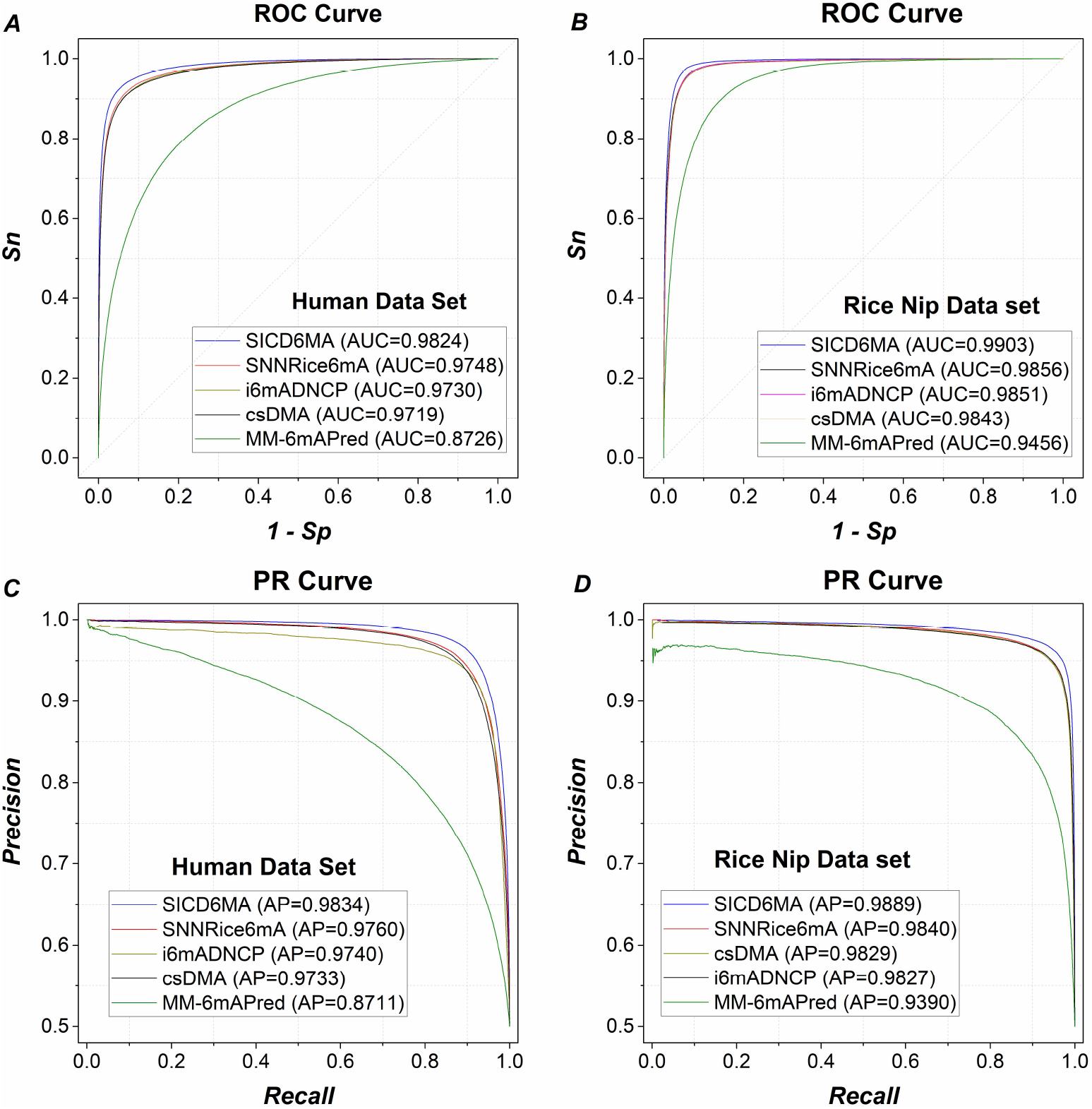
Comparisons of performance evaluation based on independent test sets. **A.** Receiver Operating Characteristic (ROC) curves of independent test evaluation based on Human data set. **B.** ROC curves of independent test evaluation based on Rice Nip data set. **C.** Precision-Recall (PR) curves of independent test evaluation based on Human data set. **D.** PR curves of independent test evaluation based on Rice Nip data set.

To investigate the impacts of several methods according to the distribution of large datasets, PR curves were plotted, as presented in **Figure 3.C** and **Figure 3.D**. Changes in PR curve were insensitive since the large datasets were balanced, and no sharp change was observed. According to these figures, the PR curves of SICD6mA were on the top of other methods. SICD6mA achieved the AP values of 0.9834 and 0.9889 in Human and Rice Nip datasets, respectively, which were the highest compared with those of other methods, indicating the highest prediction accuracy and recall of SICD6mA.Therefore, the above findings revealed that, SICD6mA attained consistent stability on the two species datasets.

To further verify the prediction performance of SICD6mA using the large dataset, these methods were compared through the accuracy and false positive rate. In this study, three thresholds, including high, medium and low, were adopted based on specificity (***Sp***) of 95%, 90%, and 80%, separately. According to **Table 1**, on human data set, SICD6mA obtained accuracy (***Acc***) of 93.66%, sensitivity (***Sn***) of 92.33%, specificity (***Sp***) of 95.00%, precision (***Pre***) of 94.86%, recall (***Rec***) of 92.33%, ***F1-score*** of 93.58% and Matthews Correlation Coefficient (***MCC***) of 87.36% at the high threshold; while ***Acc*** of 92.81%, ***Sn*** of 95.61%, ***Sp*** of 90.00%, ***Pre*** of 90.53%, ***Rec*** of 95.61%, ***F1-score*** of 93.00% and ***MCC*** of 85.75% at the medium threshold; ***Acc*** of 88.97%, ***Sn*** of 97.94%, ***Sp*** of 80.00%, ***Pre*** of 83.04%, ***Rec*** of 97.94%, ***F1-score*** of 89.88% and ***MCC*** of 79.23%at the low threshold. Thus, it was obvious that, the large datasets not only intensified the model capabilities to detect the 6mA feature signals, but also increased the ***Sn*** value, thereby reducing the false positive rate. In the meantime, the metrics of SICD6mA were apparently higher than those of other methods, showing that SICD6mA improved the prediction accuracy and precision.

**Table 1.** Performance comparison between SICD6mA and several methods on the independent test set of human genome.

| Methods | Threshold | *Acc* | *Sn* | *Sp* | *Pre* | *Rec* | *F1-score* | *MCC* | *p-value* |
| --- | --- | --- | --- | --- | --- | --- | --- | --- | --- |
| SIC6mA | High | **93.66** | **92.33** | 95.00 | **94.86** | **92.33** | **93.58** | **87.36** | - |
|  | Medium | **92.81** | **95.61** | 90.00 | **90.53** | **95.61** | **93.00** | **85.75** |  |
|  | Low | **88.97** | **97.94** | 80.00 | **83.04** | **97.94** | **89.88** | **79.23** |  |
| i6mA-DNCP | High | 91.77 | 88.52 | 95.03 | 94.68 | 88.52 | 91.49 | 83.72 | <0.1% |
|  | Medium | 91.70 | 93.28 | 90.12 | 90.42 | 93.28 | 91.83 | 83.44 |  |
|  | Low | 88.35 | 96.64 | 80.07 | 82.90 | 96.64 | 89.24 | 77.78 |  |
| MM-6mA Pred | High | 71.89 | 48.78 | 95.00 | 90.70 | 48.78 | 63.44 | 49.37 | <0.1% |
|  | Medium | 76.74 | 63.47 | 90.00 | 86.39 | 63.47 | 73.18 | 55.46 |  |
|  | Low | 79.27 | 78.53 | 80.00 | 79.70 | 78.53 | 79.11 | 58.54 |  |
| SNNRice6mA | High | 92.15 | 89.31 | 95.00 | 94.70 | 89.31 | 91.92 | 84.45 | <0.1% |
|  | Medium | 91.97 | 93.94 | 90.00 | 90.38 | 93.94 | 92.12 | 84.00 |  |
|  | Low | 88.47 | 96.93 | 80.00 | 82.90 | 96.93 | 89.37 | 78.06 |  |
| csDMA | High | 91.74 | 88.49 | 95.00 | 94.65 | 88.49 | 91.47 | 83.67 | <0.1% |
|  | Medium | 91.57 | 93.10 | 90.03 | 90.33 | 93.10 | 91.69 | 83.17 |  |
|  | Low | 88.25 | 96.43 | 80.07 | 82.87 | 96.43 | 89.13 | 77.54 |  |

Note:
1. *p-value* is the probability of the two-sample T-test between the current method and SICD6mA. 2. The threshold is the prediction score(for SICD6mA) or probability(for other methods) on independent test sets when *Sp* is equal to 95%.

These methods were compared and evaluated using the performance metrics in Rice Nip dataset. As shown in **Table 2**, SICD6mA was the optimal method, with ***Acc*** of 95.99%, ***Sn*** of 96.97%, ***Sp*** of 95.00%, ***Pre*** of 95.1%, ***Re c*** of 96.97%, ***F1-score*** of 96.02% and ***MCC*** of 91.99% at the high threshold; ***Acc*** of 94.47%, ***Sn*** of 98.94%, ***Sp*** of 90.01 %, ***Pre*** of 90.83%, ***Rec*** of 98.94%, ***F1-score*** of 94.71% and ***MCC*** of 89.31% at the medium threshold; ***Acc*** of 89.8%, ***Sn*** of 99.58%, ***Sp*** of 80.02%, ***Pre*** of 83.29%, ***Rec*** of 99.58%, ***F1-score*** of 90.71% and ***MCC*** of 81.17% at the low threshold. Besides, the metrics of SICD6mA were much greater than those of other methods, with the only exception of ***Pre*** at the low threshold. Such results demonstrated that, SICD6mAachieved higher accuracy and lower false positive rate than the existing methods with regard to Rice Nip dataset. Consequently, SICD6Ma was associated with high prediction quality and stability.

**Table 2.**
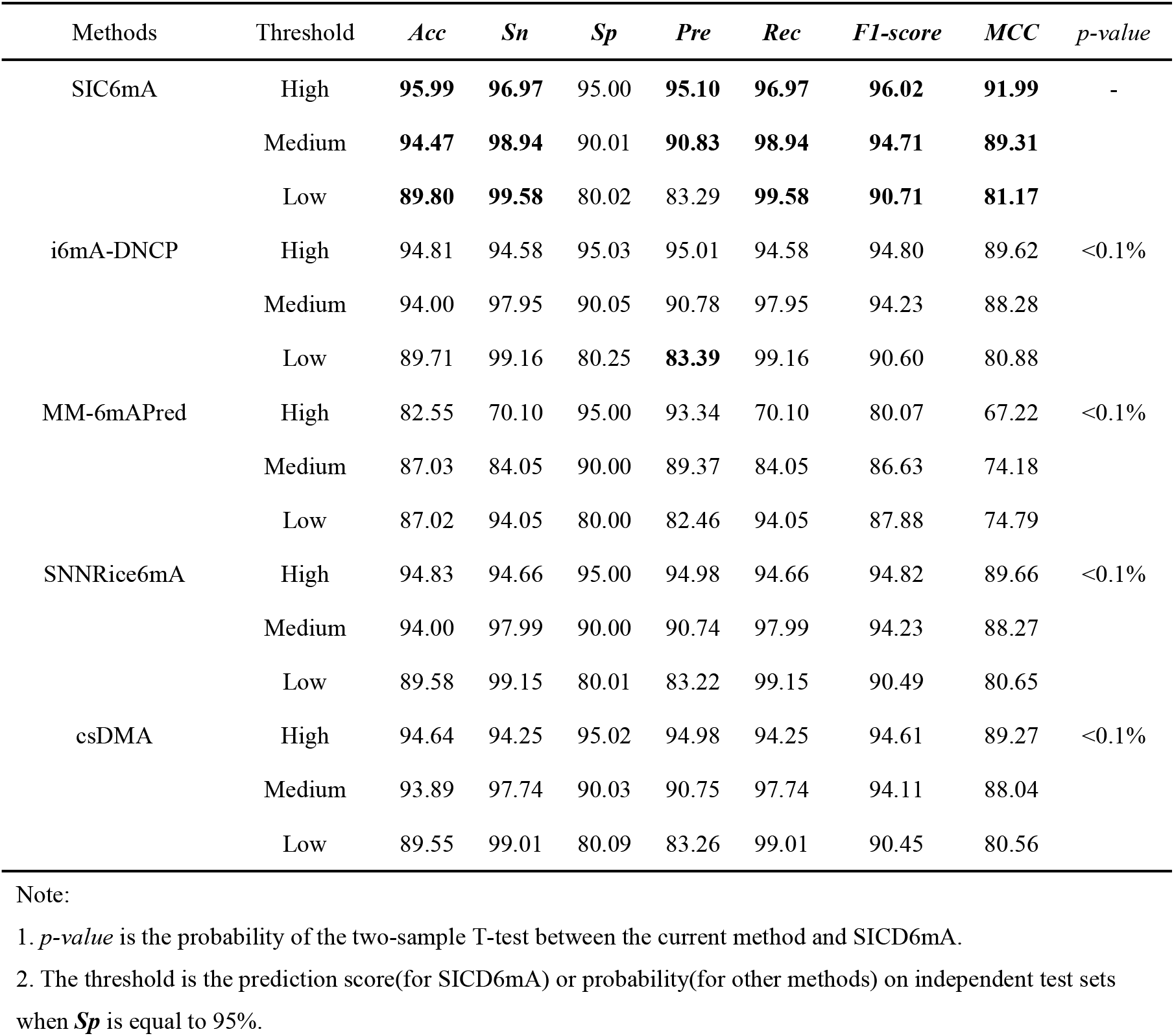
Performance comparison between SICD6mA and several methods on the independent test set of Rice Nip genome

i6mADNCP, csDMA, MM-6mAPred and other similar methods have combined the chemistry and frequency of nucleotides, and they classify sequences through traditional machine learning. However, SICD6mA is a deep learning network, with no manual design of feature extraction method. In this study, the AUC values, AP values and several performance metrics of SICD6mA were superior to those of other methods, which indicated that it was advantageous to use SICD6mA for identifying the 6mA site.

### Comparison of performance in identifying 6mA sites across species

To prove the ability of SICD6mA in sensitively identifying 6mA sites in other species, the successful prediction rates of several methods were tested using the cross-species positive datasets. To simulate the processing scene of experimental data, the CD-HIT tool was not adopted to remove the redundant positive sequences. The above models that had been trained on training datasets were employed in every method to predict the positive sites. Typically, the sequence that predicted a higher score or probability than the threshold was deemed as the6mA site sequence of successful identification; otherwise it was regarded as mis-classification. Notably, the threshold was the predicted score (for SICD6mA) or probability (for other methods) of the above independent test set when ***Sp*** was 95%.

Based on the Human dataset, the successful prediction rates of ***Arabidopsis thaliana**, **Drosophila**, **Fragaria vesca**, **Rice indica**, **Rosa chinensis**, **Tolypocladium*** and ***yeast*** were as high as 82.02%, 82.13%, 84.50%, 84.32%, 84.64%, 49.20% and 57.06%, by SICD6mA, respectively (**Table S1**). Apparently, the results obtained for ***Drosophila***, ***Tolypocladium***, ***yeast***, ***Fragaria vesca*** and ***Rosa chinensis*** were notably higher than those obtained by i6mA-DNCP, MM-6mAPred, SNNRice6mA and csDMA. Meanwhile, results for ***Arabidopsis thaliana*** and ***Rice indica*** were close to those obtained by SNNRice6mA and csDMA respectively. The mean successful prediction rate achieved 74.84%, which was higher than those of the other methods. Using the Rice Nip dataset, the successful prediction rates for ***Arabidopsis thaliana***, ***Drosophila***, ***Fragaria vesca***, ***Rice indica***, ***Rosa chinensis***, ***Tolypocladium*** and ***yeast*** reached 75.62%, 68.18%, 94.72%, 97.02%, 92.48%, 41.56% and 39.40%, respectively, by SICD6mA (**Table S2**). All results and the mean successful prediction rate (72.71%) were notably higher than those obtained by i6mA-DNCP, MM-6mAPred, SNNRice6mA and csDMA. It was figured out after integrating tables S3 and S4 that, each method attained the highest successful prediction rate (**Table 3**). The successful prediction rates achieved by SICD6mA were 82.13%, 94.72%, 97.02%, 92.48%, 49.20% and 57.06% for ***Drosophila***, ***Fragaria vesca***, ***Rice indica***, ***Rosa chinensis***, ***Tolypocladium*** and ***yeast***, separately, which were higher than those attained by other methods. The successful prediction rate (82.02%) of ***Arabidopsis thaliana*** attained by SICD6mA was close to that acquired by SNNRice6mA, and the mean successful prediction rate (79.26%) of SICD6mA was the highest compared with those of i6mA-DNCP, MM-6mAPred, SNNRice6mA and csDMA.

**Table 3.** Performance comparison for identifying 6mA sites based on human and Rice Nip data sets.

| Genome | SIC6mA | i6mA-DNCP | MM-6mAPred | SNNRice6mA | csDMA |
| --- | --- | --- | --- | --- | --- |
| *Arabidopsis thaliana* | 82.02 | 82.70 | 54.95 | **82.71** | 81.37 |
| *Drosophila* | **82.13** | 75.64 | 49.43 | 78.04 | 77.28 |
| *Fragaria vesca* | **94.72** | 92.58 | 75.71 | 93.15 | 92.10 |
| *Rice indica* | **97.02** | 95.74 | 71.27 | 94.67 | 95.56 |
| *Rosa chinensis* | **92.48** | 90.34 | 71.03 | 90.60 | 89.81 |
| *Tolypocladium* | **49.20** | 43.82 | 23.20 | 46.97 | 47.18 |
| *yeast* | **57.06** | 50.39 | 32.78 | 54.27 | 51.10 |
| Mean of Success Rates | **79.26** | 75.89 | 54.05 | 77.20 | 76.34 |

The successful prediction rates for ***Fragaria vesca***, ***Rice indica*** and ***Rosa chinensis*** attained by SICD6mA were much higher than those by other methods using the Rice Nip dataset. In the meantime, the successful prediction rates of ***Drosophila***, ***Tolypocladium*** and ***yeast*** achieved by SICD6mA are significantly higher than those obtained by other methods using the Human dataset, indicating that SICm6A was more accurate in some genomes. Therefore, SICm6A will be a very useful tool to analyze 6mA across the genome-wide.

### Application of the program

The computer memory used to predict the 6mA site is quite large, which consumes lots of calculation time. For the convenience of most experimental scientists, a 6mA site prediction tool, SICD6mA (**Figure 4**), has been developed in this study, which provides two prediction modes of Human and Rice Nip. It allows to predict the 6mA sites for human and rice species, respectively. If the user inputs the sequences and presses the button “submit”, the tool will filter the sites with “A” at the center and calculate the corresponding scores. Written using python, this tool can rapidly identify 6mA sites based on the CPU or GPU platforms.

**Figure 4.**
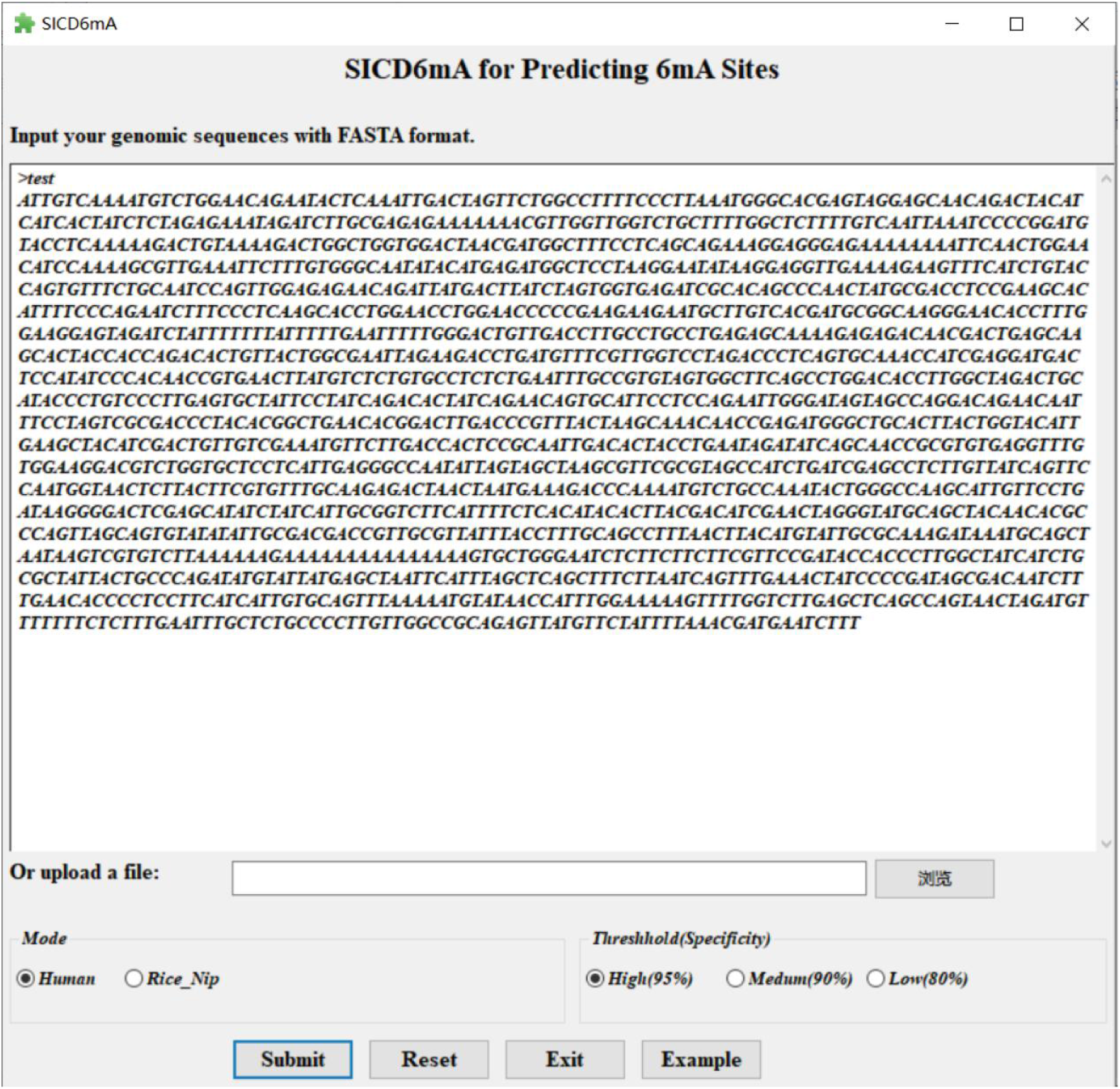
Main interface of SICD6mA prediction tool.

## Discussion

At present, prediction methods of the 6mA sites are mainly aimed at the rice genome, and it is not yet possible to accurately predict the 6mA sites in mammals such as humans. Therefore, based on deep learning, we built a prediction method for 6mA sites, SICDm6A. SICDm6A used a very efficient 3-mer ID coding, which is the serial number of three consecutive nucleic acids. Compared with traditional coding, not only SICDm6A was simple and intuitive, but also we did not have to design complex methods for feature extraction. SICDm6A had a GRU network with a memory mechanism, which could learn 3-mer ID coding and extract features of 6mA sites. Transpose feature map was placed in the middle of the first two GRU layers, and the latter GRU layer was a simplified GRU layer specifically designed in the present study. This novel method was similar to filter the image horizontally and vertically, extracting the features of the 6mA site through a specific memory mechanism. Thus, through improved methods, SICDm6A cleverly solved the problem of cross-species prediction by deep learning, so it could quickly capture 6mA site features, and gained higher prediction performance than other tools.

Previously, we found that several existing tools had the problem of high false positives. When the A sites on a sequence were predicted, many negative sites A were predicted as positive 6mA sites. After carrying out the analysis, we found the method of building the negative sequence set in their benchmark datasets was problematic. Additionally, these benchmark datasets also selected negative short sequences based on the 6mA motif, which prevented prediction tools from effectively learning the negative features of non-motif A sites. The human and rice datasets that we built earlier also used the 6mA motif to create negative sequences. All methods trained on the datasets also had a high false positive rate. Later, we switched the method of selecting negative sequences through the A site. All methods effectively avoided the problem of high false positive rate. At the same time, the prediction performance of existing tools was significantly improved on our benchmark data sets. In future research, we hope to build a 6mA benchmark dataset of prokaryotes to predict the 6mA site of species such as bacteria.

## Conclusions

N6-methyladenine (6mA) is closely linked to a series of biological processes, but the relationship of the uneven distribution of these sites with its biological functions remains largely unknown so far. To better understand the biological function of 6mA, the 6mA sites in the genome-wide have been detected experimentally. Nevertheless, the experimental techniques are time-consuming and expensive, making it necessary to develop a fast calculation method. In this study, a deep learning model SICD6mA is proposed to identify the 6mA sites in eukaryotic genomes, such as human and rice. SICD6mA encodes the sequence using the order numbers of three consecutive nucleic acids *(3-mer* ID), then extracts the information of interest and identifies the 6mA sites by the Gated Recurrent Units (GRU) networks based on the memory mechanism. In addition, the large benchmark datasets of human and rice are constructed from the initially reported 6mA sites, and the SICD6mA model is subsequently trained based on these datasets. According to a series of performance evaluations, SICD6mA attains stable performance on various species datasets, and obtains high accuracy and sensitivity. Besides, the AUC values of human and rice independent test sets reach 0.9824 and 0.9903, respectively, and the cross-species successful prediction rates are superior to those of the existing methods, thus confirming the effectiveness of the 3-*mer* ID features and the deep memory networks. More importantly, SICD6mA is also an open-source tool that can quickly identify the 6mA sites across species by supporting GPU-accelerated operations. Therefore, SICD6mA will be a useful computational tool to identify the 6mA sites in eukaryotic genomes like human and rice.

## Methods

### Dataset collection and generation

In this study, the benchmark datasets(**Table S3**) of human and rice were adopted for model training and performance evaluation, and the cross-species prediction dataset(**Table S4**) was utilized for comparing the identification ability of 6mA sites. In the datasets, the sequence length was 41nt, and A was located in the middle of the sequence. If the left side of A was shorter than 20, some gap symbols “-”were added to the front, so that the sequence length was equal to 41; similarly, gap symbols were also added to the right if its right length was shorter. Thereafter, every benchmark dataset was randomly split into a training set, and an independent test set at a ratio of 8:2, and such ratios between positive and negative sequences were maintained consistent on the training and independent sets, respectively.

The single-species benchmark datasets (**Table S3**) included two types, namely, Human (***Homo sapiens)*** and Rice Nip (***Nipponbare japonica**),* which were created in this work. The 6mA positive sequences of Human were derived from 6mA-IP-sequencing[10] that was deposited into the Gene Expression Omnibus (accession number GEO: GSE104475), and then the redundancy was removed by CD-HIT-EST[26] using the threshold of 0.8. Finally, altogether 614,856 sequences were remained. At the same time, the human negative sequences were extracted from the GRCh38/hg38 genome (http://hgdownload.soe.ucsc.edu/). There are over 100 million bases A in the human genome. Nonetheless, human negative sequences were generated due to the insufficient computing resources, as shown below, the short sequences without duplication were only searched from the forward and reverse strands in this genome based on the base “A” in the middle and 41nt length. In addition, short sequences identical to positive sequences were deleted, and a total of 770,690 sequences were randomly selected. CD-HIT-EST was also used to remove the redundancy, and 614,856 sequences were retained randomly. On the other hand, the positive sequences of Rice Nip were downloaded from the eRice[27] websites (http://www.elabcaas.cn/rice/downloads.html). CD-HIT-EST was applied in removing the redundancy of positive sequences, and eventually 380,650 positive sequences were remained. In the meantime, the negative sequences were extracted from the (https://rgp.dna.affrc.go.jp/E/IRGSP/Build5/build5.html) of ***Nipponbare japonica*** genome[28]. Altogether 761,300 short sequences distinct from positive sequences were randomly selected from the forward and reverse strands of the genome according to the base “A” in the middle and the length of 41nt. Thereafter, CD-HIT-EST was adopted for removing the redundancy, 662,710 balanced sequences were obtained, and 380,650 negative sequences were randomly remained.

The cross-species prediction dataset was established based on 6mA positive sequences of several eukaryotic genomes (**Table S4**), including ***Arabidopsis thaliana, Fragaria vesca, Rosa chinensis, Rice indica, Drosophila, Tolypocladium*** and ***yeast***. Among them, the positive sequences of ***Drosophila, Tolypocladium*** and ***yeast*** were downloaded from the MethSMRT[29] database (http://sysbio.sysu.edu.cn/methsmrt/), and the sequences described as “6mA”were deemed as the 6mA positive sequences. The positive sequences of ***Rice indica*** were derived from the eRice websites (http://www.elabcaas.cn/rice/downloads.html), while those of ***Arabidopsis thaliana***[9] were collected from the NCBI Gene Expression Omnibus (GEO) with accession number GSE81596 (GSM2157793), and those of ***Fragaria vesca***[30], and ***Rosa chinensis***[31] were obtained from the MDR database (http://mdr.xieslab.org/).

### Sequence Representation

#### Nucleic Acid Encoding

The different chemical properties of four nucleic acids (A,G) and (T,C) are that, they have one and two rings, respectively; besides, each (A,C)and(T,G)contain the aminoandketogroups; whereas (A,T)and(C,G) possess twoandthree hydrogenbonds, respectively. These properties can be computed as follow[20]:

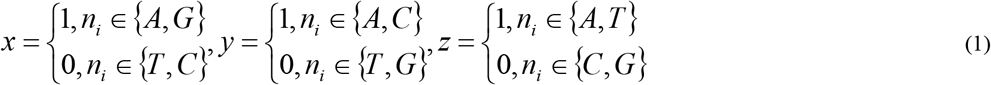

Thus, a nucleic acid is represented as a one-dimensional vector (*x, y, z*); in other words, A, G, C and T to (1,1,1), (1,0,0), (0,1,0)and (0,0,1), respectively. This suggests that, chars A, G, C and T have the sort encoding of 1, 2, 3, and 4, respectively, if their vectors are converted the numbers of 111, 100, 10 and 1, separately. In addition, the encoding of gap symbol “N” or “-”is 0in a sequence.

#### *3-Mer* ID Feature

In this study, *3-Mer* ID represented the three chars from nucleic acids or gap symbol, which was encoded as an integer by combining the encoding sequentially; for example, TAG -> 412, CGA -> 321, and GA-->210 (**Figure 1.A**). A sequence of *n* bits might be converted to *n* integers through*3-Mer* ID encoding. To make the converted length consistent with *n*, two gap symbols “-”are added after the last nucleic acid at first. Next, every three consecutive chars are extracted from left to right in turn in the sequence, after that, all *3-Mer* chars are encoded as *3-mer* ID one by one. Therefore, the sequence changes to an integer vector, namely, the *3-Mer* ID feature.

### Construction of the SICD6mA model

Memory network is the model that describes the long-term dependencies of sequential data, and the memory components are used to save scene information, achieve long-term memory^[32]^ and extract important features. The Recurrent Neural Network (RNN), long short-term memory (LSTM) and Gated Recurrent Units (GRU) layers of deep learning are associated with certain memory mechanisms. Combined with other deep learning layers, a variety of deep memory networks can be constructed to deal with the sequential data such as the biological sequences.

#### Word-Vector Embedding Layer

It is found that the relationship between variables can be expressed quantitatively using a numerical matrix. Typically, LSTM is a recursive neural network for learning the long-term dependency, namely, a numerical description matrix between nucleic acids in a sequence. GRU is the simplified version of LSTM, which is sometimes superior and faster. Nonetheless, GRU is unable to directly handle the site coordinates between nucleic acids, and it is only capable of reading numerical matrices that describe the original features. Thus, a layer called the word vector embedding[33] is required to convert the site information into the original feature matrix. In this study, after the nucleic acid sequence was transformed into a *3-Mer* ID feature with a length of *n* nt, each site corresponded to a *3-Mer* ID, which was an integer (**Figure 1.B**). Then, the word vector embedding layer functioned to map the integers at each site into a *m*-dimensional numerical vector. In this paper, *m* was equivalent to 125. After converting a nucleic acid sequence with *n* nt into a numerical matrix of *nxm,* important features were discovered from the GRU layer by learning the numerical matrix.

#### GRU Layer

It is more difficult for the recurrent neural networks to capture the dependency of nucleic acids with huge separation distances in the sequences, which is ascribed to the problem of gradient attenuation during training[34]. In this study, the proposed GRU was able to better capture the sequence dependency in the long distance (**Figure 1.B)**. The learning process of GRU conformed to the decay memory mechanism of human, to be specific, the fuzzy memory of past things and clear memory of recent things. This attenuation effect was determined by the product of the attenuation coefficient and the memory value. In addition, the sigmoid function was employed to generate the attenuation coefficient, and the memory value was obtained via the tanh function. This learning mechanism was exhibited in equation (2) – (5).

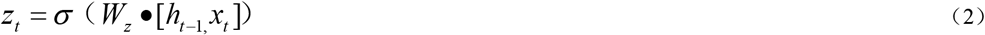

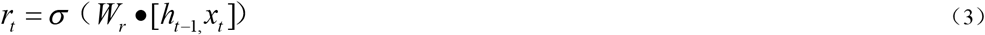

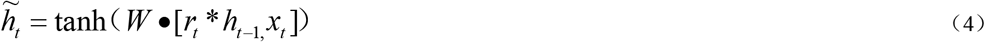

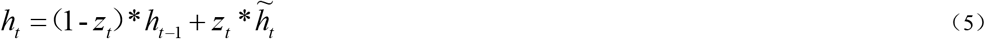

where *x_t_* represents the current features for learning, *h*_*t*-1_ stands for the features remembered after previous learning, 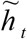 is equivalent to the first impression generated before learning, *z_t_* and *r_t_* indicate the attenuation coefficients, *h_t_* suggests the features to be remembered after learning, • and * represent the product and dot operations of the matrix, whereas *W_z_*, *W_r_, W* are the model parameters representing the network ability to learn features, and these parameters are automatically fitted through model training. GRU achieves two attenuation effects, namely, when the first impression and final memory are generated, respectively. Of them, the first impression is affected by previous memory, while the final memory is composed of the sum of decayed values from the previous memory and the first impression memory.

The Bi-directional Gated Recurrent Units (BGRU) network suggests that, each training sequence is made up of two recurrent neural networks (forward and backward), which are connected to an output layer. UGRU is short for the non Bi-directional Gated Recurrent Units, which is the GRU.

#### SGRU Layer

As found from the above discussion, the decay memory mechanism of GRU is substantially disturbed by the previous memory, which is not conducive to discovering the new features that do not change significantly. Consequently, a memory layer called the Simple Gated Recurrent Units (SGRU) layer, which possesses two activation functions of tanh and sigmoid, was specifically designed in this study (**Figure 1.B**). Among them, the tanh function generates candidate memories, while the sigmoid function builds attenuation coefficients, and these two functions produce the effect of remembering important information and forgetting the unimportant one. SGRU synthesizes the valid memory using the results of these two functions; in this way, it extracts the abstract features by forgetting the unimportant features.

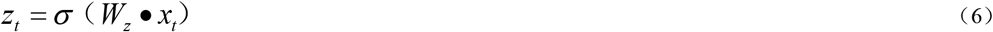

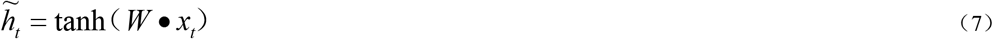

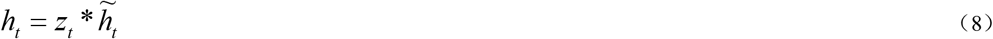

where *x_t_* represents the current features for learning, 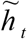 is equivalent to the first impression generated before learning, *h_t_* stands for the features to be remembered after learning, while • and * suggest the product and dot operations of matrix, respectively. Notably, SGRU has the one-time attenuation effect when the final memory is generated, its first impression remains unaffected by the previous memory, and the final memory stands for the attenuation value of the first impression memory. This memory network is able to discover the subtle differences between sequence features.

#### Other Layers

The LeakyReLU activation function divides all negative values by a constant of >1. Among them, the dropout layer means that all values less than a certain threshold are all set to 0, and this layer prevents overfitting by dropping operation when too few sample data are used for model training. During the process of training, BatchNorm1d normalizes data by batch, which also prevents overfitting. Moreover, the fully connected layer is equivalent to a “classifier” in the network. The log_softmax function generates the probability value from the numerical features learned, and later converts them into log values, which are equivalent to the scoring values.

#### Parameters of SICD6mA

SICD6mA could accelerate training on GPU platform using Adam optimizer and learning rate of 0.0001. It adopted the loss function of Cross-Entropy Loss, and the num_embeddings and embedding_dim of the word-vector embedding layer were 1024 and 125, separately. At the same time, the number of BGRU layers was 2, input_size was 125, hidden_size was 125, dropout was 0.5, batch_first was “True”. and the bidirectional was equal to “True”. On the other hand, the number of UGRU layers was 2, input_size was 51, hidden_size was 128, dropout was 0.5, batch_first was “True”. and the bidirectional was equal to “False”. As for SGRU. the parameters were 125 and 250, respectively, and the fully connected layers were FC1: Output=16 and FC2: Output=2. Besides, the three layers of LeakyReLU (negative_slope=0.2), BatchNorm1d (momentum=0.5) and Dropout (p=0.25) were in the middle of FC1 and FC2.

### Performance evaluation

The receiver operating characteristic (ROC) curve was adopted to evaluate the model performance. Typically, the area under the ROC curve (AUC) is an important indicator for binary classification, which is a float value between 0 and 1, and a AUC value closer to 1 suggests the better model performance. In addition, the PR curve is the precision *vs* recall curve, in which recall serves as the abscissa axis, while precision as the ordinate axis, and a larger area under the PR curve (namely, the AP value) is indicative of the higher accuracy.

In this study, the prediction performance of classifiers was evaluated by five metrics: accuracy (***Acc***), sensitivity (***Sn***), specificity (***Sp***), precision (***Pre***), recall (***Rec***), ***F1-score*** and Matthews Correlation Coefficient (***MCC***), as defined below:

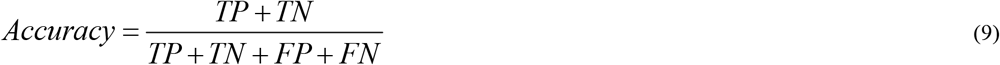

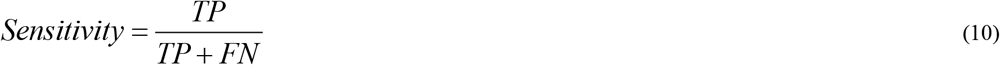

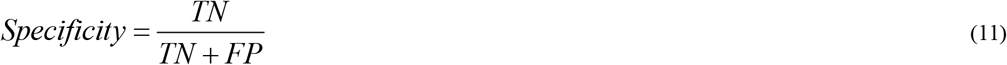

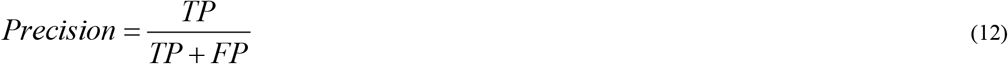

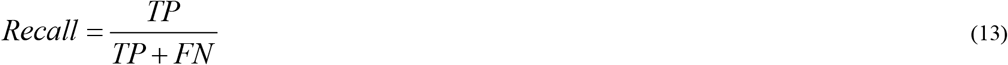

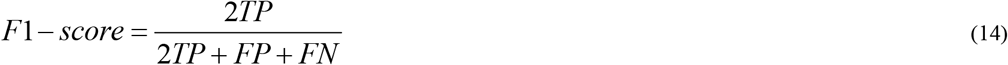

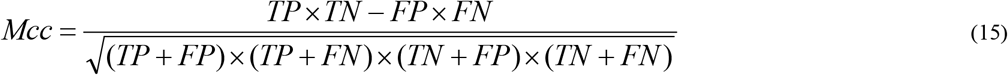

Where TP (true positive) is the correctly classified number of positive samples, TN (true negative) represents the correctly predicted number of negative samples, FP (false positive) stands for the incorrectly classified number of negative samples, and FN (false negative) is the error count in predicting the positive samples.

## List of abbreviations

6mA: DNA N6-methyladenine
SMRT: single-molecule real-time sequencing
SVM: support vector machines
SICD6Ma: Identifying 6mA Sites using Deep Memory Network
Rice Nip data set: 6mA sites data set of ***Nipponbare japonica*** genome
Human data set: 6mA sites data set of ***Homo sapiens*** genome
iDNA6mA-PseKNC: Identifying DNA N6-methyladenosine sites by incorporating nucleotide physicochemical properties into PseKNC
i6mA-Pred: identifying DNA N6-methyladenine sites in the rice genome
MM-6mAPred: identifying DNA N6-methyladenine sites based on Markov model
SDM6A: a web-based integrative machine-learning framework for predicting 6mA sites in the rice genome
csDMA: an improved bioinformatics tool for identifying DNA 6 mA modifications via Chou’s 5-step rule
i6mA-DNCP: Computational Identification of DNA N6-Methyladenine Sites in the Rice Genome Using Optimized Dinucleotide-Based Features
SNNRice6mA: a deep learning method for predicting DNA N6-methyladenine sites in rice genome
eRice: a refined epigenomic platform for japonica and indica riceg
MethSMRT: an integrative database for DNA N6-methyladenine and N4-methylcytosine generated by single-molecular real-time sequencin
6mA-rice-Lv: a benchmark dataset of rice genome
6mA-rice-Chen: a benchmark dataset of rice genome
3-*mer* ID: the serial numbers of three consecutive nucleic acids. It was encoded a sequence in this study.
GRU: the Gated Recurrent Units
UGRU: non Bi-directional Gated Recurrent Units
BGRU: Bi-directional Gated Recurrent Units
SGRU: Simple Gated Recurrent Units
RNN: Recurrent Neural Network
LSTM: long short-term memory
GPU: Graphic Processing Unit
CPU: Central Processing Unit
ROC: receiver operating characteristic
AUC: area under the curve
CD-HIT: A package can perform various jobs like clustering a protein database, clustering a DNA/RNA database, comparing two databases (protein or DNA/RNA), and generating protein families.
CD-HIT-EST: One tool of CD-HIT. It clusters a nucleotide sequences that meet a similarity threshold, usually a sequence identity.
GEO: Gene Expression Omnibus
*Acc*: accuracy
*Sn*: sensitivity
*Sp*: specificity
*Pre*: precision
*Rec*: recall
*F1-score*: harmonized average of precision and recall
*MCC*: Matthews Correlation Coefficient

## Declarations

### Ethics approval and consent to participate

Not applicable.

### Consent for publication

Not applicable.

### Availability of data and materials

The datasets supporting the conclusions of this article are available at https://github.com/lwzyb/SICD6mA.

### Author’s contribution

Wenzhong Liu and Hualan Li conceived the study. Wenzhong Liu designed the algorithms and the software. Hualan Li tested the software. Wenzhong Liu wrote the manuscript. The final version of the manuscript is approved by all authors.

## Acknowledgements

Not applicable.

## Supplementary material

**Table S1.** Performance comparison for identifying 6mA sites based on human data set

| Genome | SIC6mA | i6mA-DNCP | MM-6mAPred | SNNRice6mA | csDMA |
| --- | --- | --- | --- | --- | --- |
| *Arabidopsis thaliana* | 82.02 | 82.70 | 54.95 | **82.71** | 81.37 |
| *Drosophila* | **82.13** | 75.64 | 49.43 | 78.04 | 77.28 |
| *Fragaria vesca* | **84.50** | 81.44 | 75.71 | 83.39 | 83.15 |
| *Rice indica* | 84.32 | 82.21 | 71.27 | 83.39 | **84.54** |
| *Rosa chinensis* | **84.64** | 82.77 | 71.03 | 83.68 | 84.49 |
| *Tolypocladium* | **49.20** | 43.82 | 23.20 | 46.97 | 47.18 |
| *yeast* | **57.06** | 50.39 | 32.78 | 54.27 | 51.10 |
| Mean of Prediction Success Rates | **74.84** | 71.28 | 54.05 | 73.21 | 72.73 |

**Table S2.** Performance comparison for identifying 6mA sites based on Rice Nip data set

| Genome | SIC6mA | i6mA-DNCP | MM-6mAPred | SNNRice6mA | csDMA |
| --- | --- | --- | --- | --- | --- |
| *Arabidopsis thaliana* | **75.62** | 70.57 | 52.29 | 72.22 | 68.77 |
| *Drosophila* | **68.18** | 57.31 | 47.10 | 63.99 | 57.50 |
| *Fragaria vesca* | **94.72** | 92.58 | 73.32 | 93.15 | 92.10 |
| *Rice indica* | **97.02** | 95.74 | 68.55 | 94.67 | 95.56 |
| *Rosa chinensis* | **92.48** | 90.34 | 68.58 | 90.60 | 89.81 |
| *Tolypocladium* | **41.56** | 36.81 | 21.52 | 39.53 | 37.59 |
| *yeast* | **39.40** | 29.73 | 30.90 | 35.69 | 29.60 |
| Mean of Prediction Success Rates | **72.71** | 67.58 | 51.75 | 69.98 | 67.28 |

**Table S3.** Summary of the training and independent test sets

| Name of data sets | Content | Number of<br>Training sets | Number of<br>Independent test sets | Total Number |
| --- | --- | --- | --- | --- |
| **Human** | Sum. | **983,770** | **245,942** | **1,229,712** |
| includes: | Positive | 491,885 | 122,971 | 614,856 |
|  | Negative | 491,885 | 122,971 | 614,856 |
| **Rice Nip** | Sum. | **609,040** | **152,260** | **761,300** |
| includes: | Positive | 304,520 | 76,130 | 380,650 |
|  | Negative | 304,520 | 76,130 | 380,650 |

**Table S4.** Summary of the cross species prediction set

| Genome | Positive Number |
| --- | --- |
| *Arabidopsis thaliana* | 38,052 |
| *Drosophila* | 163,493 |
| *Fragaria vesca* | 26,514 |
| *Rice indica* | 776,858 |
| *Rosa chinensis* | 14,677 |
| *Tolypocladium* | 3,453 |
| *yeast* | 4,094 |

